# Non-Genetic Mechanisms of Fractional Resistance to Abemaciclib in Dedifferentiated Liposarcoma

**DOI:** 10.64898/2026.05.22.727236

**Authors:** Lauryn E. Bailey, Samuel C. Wolff, Tarek M. Zikry, Garrett A. Sessions, Austin A. Whitman, Elizaveta K. Titerina, Hayden Raish, Joal D. Beane, Jeremy E. Purvis, Philip M. Spanheimer

**Author notes:** **Correspondence should be addressed to:** Philip M. Spanheimer MD, Department of Surgery and Department of Genetics Lineberger Comprehensive Cancer, University of North Carolina at Chapel Hill, 170 Manning Drive, Physicians Office Building Suite 1150 Chapel Hill, NC 27599-7213 USA; Jeremy E. Purvis, Ph.D., Professor and Associate Chair for Education Department of Genetics, Lineberger Comprehensive Cancer Center UNC Computational Medicine Program University of North Carolina at Chapel Hill, 11018C Mary Ellen Jones Building, 116 Manning Drive Chapel Hill, NC 27599-7488. Lauryn E. Bailey and Samuel C. Wolff contributed equally to this work.

## Abstract

Dedifferentiated liposarcoma is a rare mesenchymal malignancy driven by amplification of chromosome 12q13-15, which includes the oncogenes CDK4 and MDM2. CDK4 amplification provides a rationale for targeted therapy with CDK4/6 inhibitors, and abemaciclib has shown the most durable activity reported to date in this disease. Clinical responses, however, are incomplete and often transient, and the cellular features that allow tumor cells to continue proliferating during treatment are not well understood. To address this gap, we performed multiplexed single-cell imaging to quantify 17 cell-cycle regulators in both dedifferentiated liposarcoma cell line Lipo246 and surgically resected primary human cells exposed to abemaciclib. Both models contained a subpopulation of cells that retained phosphorylated retinoblastoma protein, a marker of cell proliferation, at the highest abemaciclib doses. These fractionally resistant cells were defined by selective enrichment of cyclin-dependent kinase 2 (CDK2), cyclin B1, and phosphorylated ribosomal protein S6 (pS6), and showed enhanced sensitivity to the CDK2 inhibitor, tagtociclib. Together, these findings reveal nongenetic cell cycle plasticity as a mechanism of escape from CDK4/6 inhibition in dedifferentiated liposarcoma and nominate CDK2 and the PI3K-mTOR pathway as candidate targets for combination therapy.

## Introduction

Liposarcoma are rare mesenchymal malignancies arising from adipose cells and occurring in the extremities, trunk, and retroperitoneum (1, 2). Well-differentiated and dedifferentiated liposarcoma constitute a major clinical subset and are characterized by recurrent amplification of chromosome 12q13-15, which includes the oncogenes MDM2 and CDK4 (3, 4). Surgery remains the primary treatment for localized disease, whereas effective systemic therapies remain limited (2, 5). Cytotoxic chemotherapy offers limited benefit and substantial toxicity, emphasizing the need for improved biologically targeted treatment strategies (2, 6).

Amplification of CDK4, a key regulator of G1/S cell cycle progression, provides a strong rationale for the use of CDK4/6 inhibitors in well-differentiated and dedifferentiated liposarcoma (3, 7). Clinical studies of palbociclib and abemaciclib have demonstrated activity in this disease (8, 9), with abemaciclib associated with a longer median progression-free survival (33 weeks vs 18 weeks for palbociclib), and durable disease control extending beyond 2 years in a subset of patients (10). However, responses remain incomplete and often transient, indicating that liposarcoma cells are not uniformly suppressed by CDK4/6 inhibition. The cellular features associated with persistence of proliferation under treatment remain poorly defined. Understanding how liposarcoma cells adapt to CDK4/6 inhibition to allow continued proliferation and establishing a panel of biomarkers which identify adapted cells is therefore essential for developing combination strategies that enhance therapeutic efficacy.

Decades of work on chemotherapy response have shown that incomplete tumor cell killing, termed fractional killing, reflects cell-to-cell variability rather than genetic resistance (11, 12). In a previous study focused on resistance to CDK4/6 inhibitors in ER+/HER2− breast tumor cells, we defined the concept of fractional resistance as a subset of tumor cells that escape therapy-induced arrest due to nongenetic cell-to-cell heterogeneity, extending the fractional killing principle to targeted cell cycle inhibitors. We showed that fractional resistance arises from cell cycle plasticity, whereby individual cells use alternative regulatory states to maintain RB phosphorylation and continued progression through the cell cycle (13). Cell cycle progression is governed largely by protein abundance, phosphorylation, and protein complex activity; therefore, identifying unique cell cycle states requires single-cell, protein-level measurements rather than transcript-level analysis alone (13, 14).

In this study, we used multiplexed single-cell imaging to quantify cell cycle regulators in a dedifferentiated liposarcoma cell line and primary human tumor cells treated with abemaciclib. We identified a resistant tumor subpopulation that continued to proliferate under CDK4/6 inhibition. These cells showed increased CDK2 and cyclin B1 expression and occupied novel cell cycle states not observed in untreated conditions, persisting even at the highest drug doses. Co-treatment with a CDK2 inhibitor reduced resistant cells in a dose-dependent manner, revealing distinct responses across conditions. Together, these findings support a model in which cell cycle plasticity enables escape from CDK4/6 inhibition and suggest targeting complementary pathways may improve efficacy.

## Results

### PHATE visualization defines proliferative region under CDK4/6 inhibition

We performed single-cell proteomic profiling using iterative indirect immunofluorescence imaging (4i) to quantify 17 cell cycle regulators within the same individual cells in both dedifferentiated liposarcoma (DDLPS) cell line Lipo246 and primary human dedifferentiated liposarcoma cells (Figure 1, A and B; see Methods)(13,14). This approach enables high-dimensional, multiplexed quantification of cell cycle state at single-cell resolution, visualized across a matrix of cells and features (Fig. 1C).

**Figure 1.**
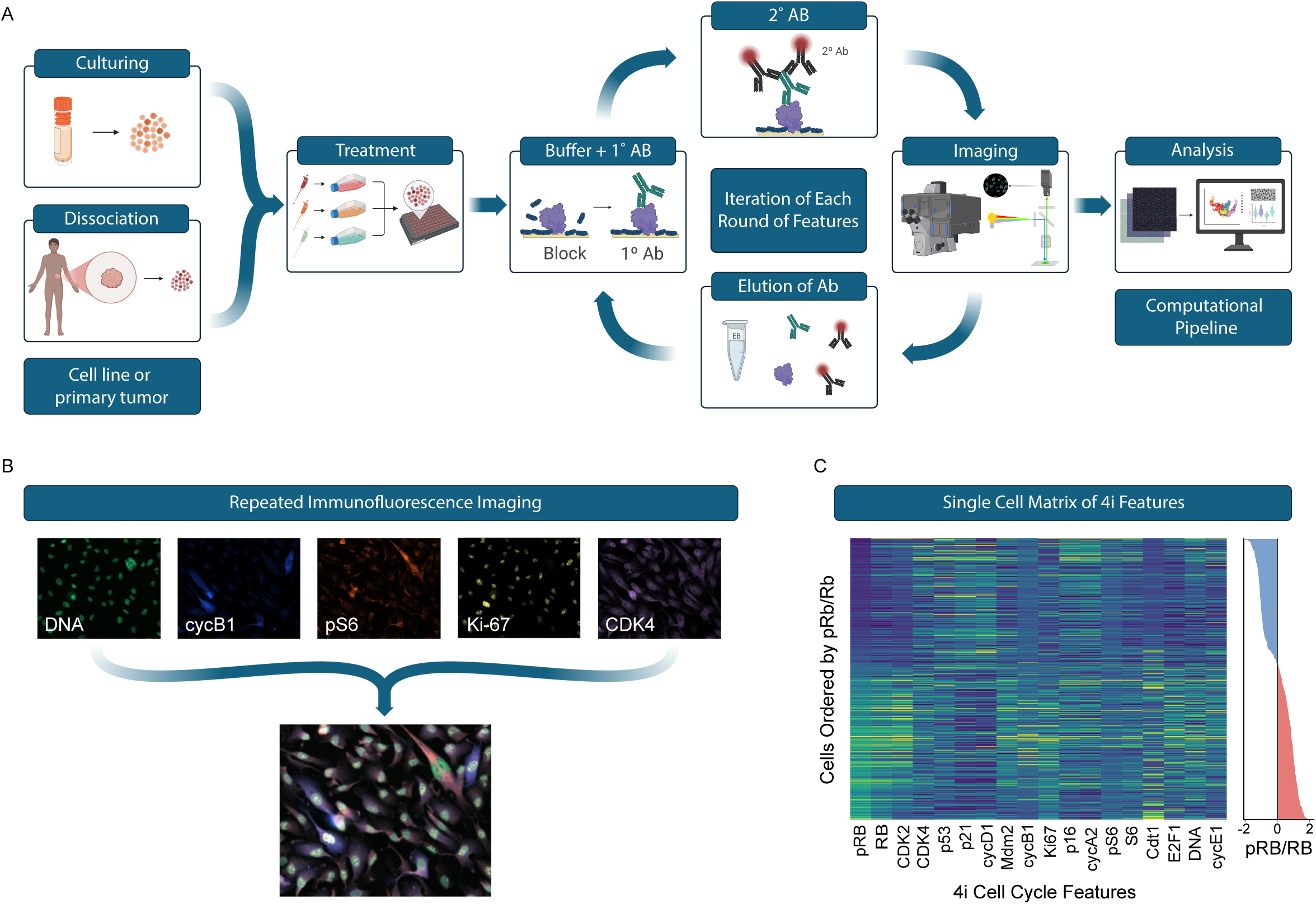
Operating room-to-lab single-cell proteomic workflow using iterative indirect immunofluorescence (4i) (A) Experimental workflow of the operating room-to-lab pipeline used for single-cell, multiplexed immunofluorescence profiling. Lipo246 cells were cultured and dedifferentiated primary tumor cells were resected and dissociated to be treated with increasing concentrations of abemaciclib for 24 h. Cells were fixed and subjected to iterative indirect immunofluorescence imaging (4i) consisting of iteration of antibody labeling (block → 1°/2° Ab → elution) and imaging. Images from all 4i rounds are processed through a computational pipeline to perform single-cell segmentation, signal quantification, and feature extraction. (B) Lipo246 and primary tumor cells were stained for 17 cell cycle regulators pRB, RB, Ki-67, CDK2, CDK4, cyclin D1 (cycD1), cyclin E1 (cycE1), Cdt1, E2F1, cyclin A2 (cycA2), cyclin B1 (cycB1), p21, Mdm2, pS6, S6, p53, p16, and integrated DNA. Representative immunofluorescence images from sequential rounds of 4i demonstrate a subset of these regulators for repeated labeling of distinct protein targets within the same cells. Nuclear DNA staining is used for segmentation, image registration, and integration across imaging rounds. (C) Single-cell matrix of 4i-measured cell cycle features. Heatmap of normalized protein expression across all quantified regulators, with individual cells ordered by increasing ratio of phosphorylated to total RB (pRB/RB), computed from 4i measurements. The corresponding single-cell pRB/RB distribution is shown at right and exhibits a bimodal structure used to define low pRB/RB (blue; nonproliferating) and high pRB/RB (pink; proliferating) cell populations. This organization reveals coordinated variation in cell cycle regulators across the pRB/RB-defined axis of cell cycle state.

In order to characterize how liposarcoma cells organize across these proteomic states under CDK4/6 inhibition, we applied Potential of Heat-diffusion for Affinity-based Transition Embedding (PHATE) to our 4i dataset (Fig. 2A)(15). PHATE transforms high-dimensional single-cell proteomic signatures into a low-dimensional embedding per-cell, with each cell represented by a single point within the PHATE structure. This low dimensional embedding preserves local and global feature relationships, such that cells positioned closer together share more similar cell cycle regulator expression profiles. The resulting embedding allowed us to visually assess how fractionally resistant cells are distributed relative to other cells across cell cycle states.

**Figure 2.**
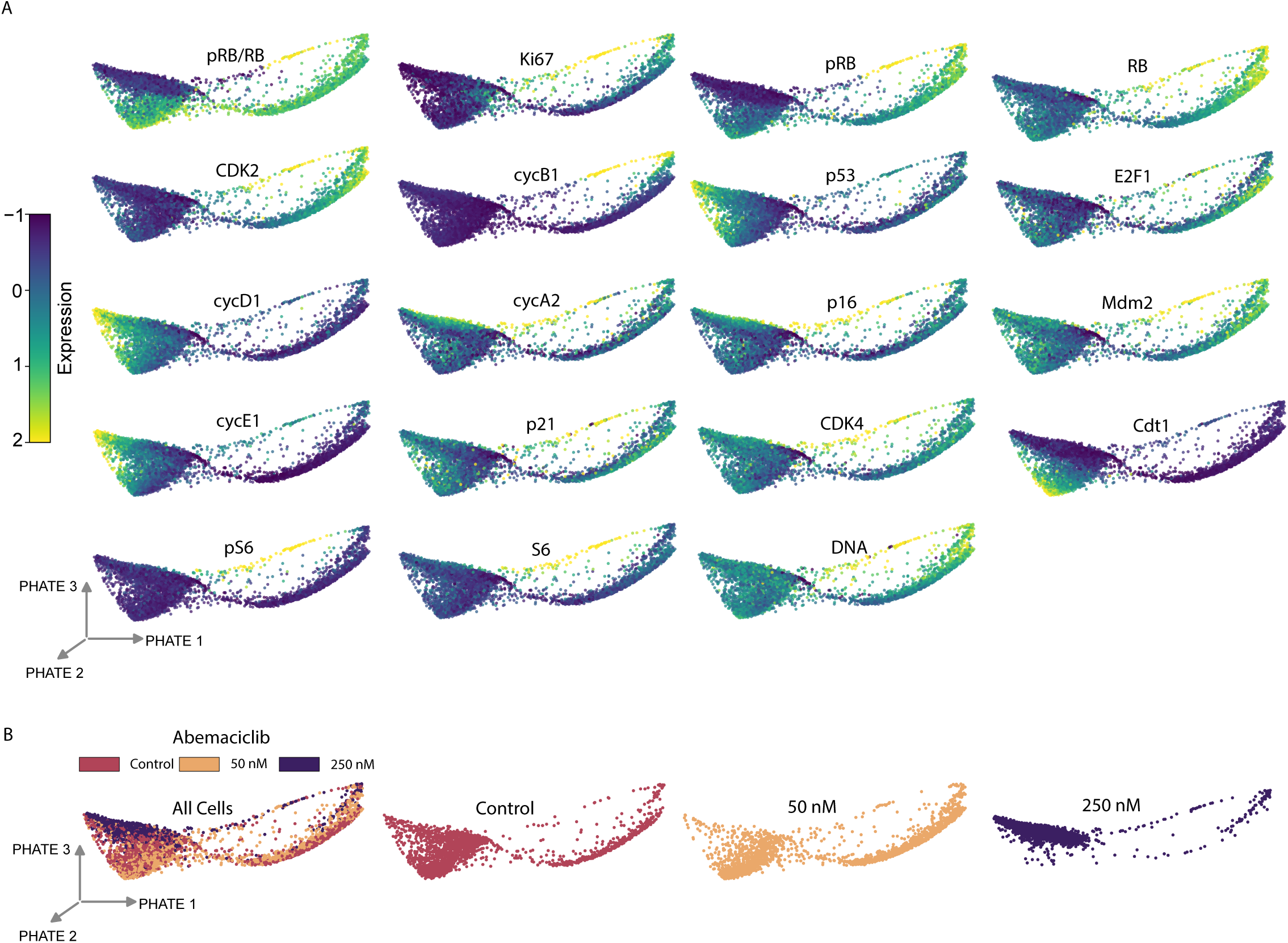
PHATE visualization of single-cell cell cycle states in Lipo246 cells under abemaciclib treatment. Single-cell protein expression profiles obtained by iterative indirect immunofluorescence imaging (4i) were down sampled and visualized using potential of heat-diffusion for affinity-based trajectory embedding (PHATE). Each point represents an individual cell, with proximity reflecting similarity in cell cycle regulator expression. Expression levels of nuclear area, phosphorylated RB (pRB), Ki-67, and additional cell cycle regulators is overlaid with color indicating relative expression (low to high). (B) Distribution of cells across treatment conditions projected onto the PHATE embedding. Cells are colored by treatment group (0, 50, and 250 nM abemaciclib) to illustrate changes in cell distribution across the embedding. Additionally, an interactive 3D PHATE viewer is hosted at https://laelise.github.io/ddlps-fractional-resistance/.

The PHATE embedding revealed a continuous curved structure consistent with progression through cell cycle states, indicating substantial variability in cell cycle progression in dedifferentiated liposarcoma states (Fig. 2A). To orient this structure, we overlaid DNA content and nuclear area, which demonstrated a transition from cells with large nuclei and high DNA content (4N) to cells with small nuclei and low DNA content (2N) along a distinct arm of the embedding, consistent with progression from pre-mitotic to recently divided states.

Overlay of pRB expression showed strong enrichment along this arm, identifying it as a proliferative region in contrast to pRB low cells outside of this region. Ki-67, a clinically established marker of proliferation, is routinely used to assess tumor proliferative fraction, with elevated Ki-67 staining associated with worse outcomes across multiple cancers (16, 17). The persistence of Ki-67 along this arm further supported classification of this region as a proliferative cell cycle state.

Having defined this proliferative region, we next examined how cells were distributed across the PHATE embedding under different treatment conditions. Cells from each treatment group (0, 50 nM, and 250 \nM abemaciclib) were colored and projected individually onto the PHATE embedding (Fig. 2B). Across these projections, increasing concentrations of abemaciclib reduced occupancy of the proliferative arm. However, even at the highest dose, a subset of cells remained localized this same region, indicating persistence of a proliferative state rather than emergence of a distinct alternative population. These observations establish that dedifferentiated liposarcoma cells occupy a continuous range of cell cycle states and that fractionally resistant cells are positioned within a persistent proliferative region under CDK4/6 inhibition.

### A subset of Lipo246 cells remains proliferative under CDK4/6 inhibitor treatment

Having identified a proliferative region, a quantitative metric to classify proliferating cells as those with an elevated ratio of phosphorylated to total RB protein (pRB/RB) was used to quantify changes in protein abundance in proliferating cells under different treatment conditions. The pRB/RB ratio exhibited a bimodal distribution across individual cells enabling an empiric cut off distinguishing proliferating (pRB/RB-high) and non-proliferating (pRB/RB-low) populations (Fig. 3A). Under control conditions, 92.3% of cells were classified as pRB/RB-high. Treatment with abemaciclib reduced the fraction of proliferating cells in a dose-dependent manner, with 85.1% remaining pRB/RB-high at 50 nM and 5.1% at 250 nM. Despite near-complete suppression of proliferation at the higher dose, a distinct pRB/RB-high population remained at 250 nM, consistent with the persistence of a fractionally resistant subset of cells. Ki-67 showed a similar dose-dependent reduction with abemaciclib treatment (*P* < 0.0001).

**Figure 3.**
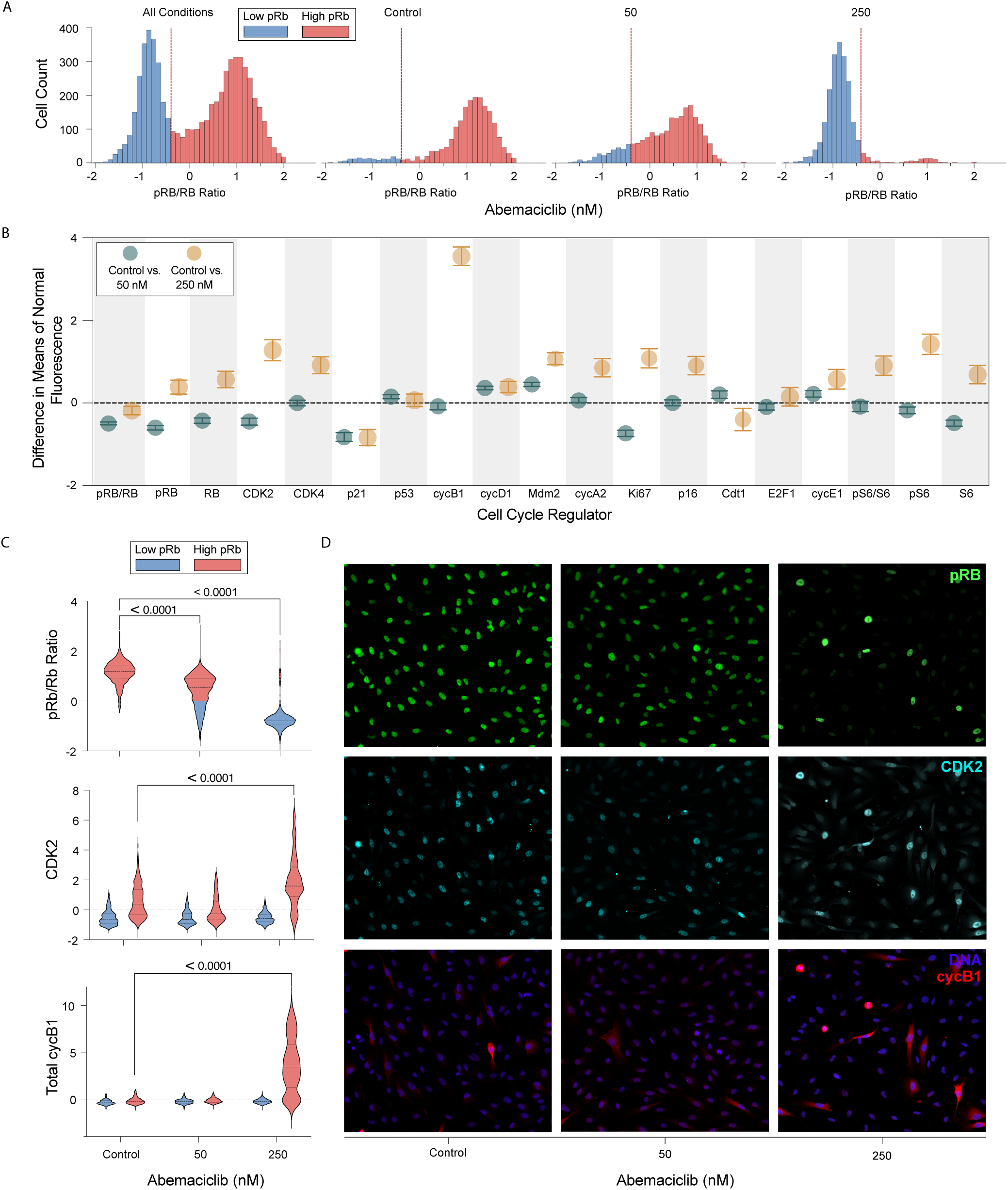
Lipo246 Cell Cycle Regulator Expression in Lipo246 Cell Cycle Regulator Expression in Fractionally Resistant Cells. (A) Distribution of the ratio of phosphorylated to total RB protein (pRB/RB) across individual Lipo246 cells under control (0 nM), 50 nM, and 250 nM abemaciclib treatment. pRB/RB exhibited a bimodal distribution, with a low pRB/RB population corresponding to hypophosphorylated RB and cell cycle arrest, and a high pRB/RB population corresponding to RB hyperphosphorylation and active proliferation. A conservative threshold (vertical red line) was defined to distinguish low pRB/RB and high pRB/RB cells and was used to classify proliferating cells for all subsequent single-cell analyses. (B) 95% CI of proliferating cells for the differences in mean expression (normalized z-scores) between either untreated cells and 50 nM abemaciclib (blue); or untreated and 250 nM abemaciclib (orange). CI overlapping with the dashed line at 0 indicate a lack of statistical significance. (C) Lipo246 cells were quantified, separated into high pRB/RB (pink) or low pRB/RB (Blue) levels. The top panel shows a dose-dependent reduction in cell cycle activity by abemaciclib as demonstrated by a reduction in pRB/RB positivity. While the middle and bottom panel shows an enrichment of CDK2 and cyclin B1 (cycB1) (respectively) in fractionally resistant cells. This enrichment is shown to be statistically significant by ANOVA analysis as seen in the corresponding violin plots. (D) Representative images of the effect of abemaciclib on Lipo246. Cells were treated for 24 hours with 0 (control), 50 nM or 250 nM abemaciclib. Cells were labeled for phosphorylated RB (pRB) which is a marker for actively cycling cells (top panel), cyclin dependent kinase 2 (CDK2)(middle panel) or DAPI for DNA and cycB1 (bottom panel). DAPI staining in the bottom panels shows roughly equal amounts of cells in each drug condition. We can visually observe the quantified depletion of pRB over increasing doses while a fraction of high pRB/RB cells persisting at high drug concentration with of upregulated CDK2 and cycB1.

### CDK2 and cyclin B1 are selectively enriched within the proliferative region under treatment

Having identified a persistent proliferative region and established a quantitative definition of proliferating cells, we next asked which cell cycle regulators were specifically associated with fractionally resistant cells under treatment. Integration of protein expression with PHATE embedding revealed that CDK2 and cyclin B1 were selectively enriched along the proliferative region, with comparatively low expression outside of this region.

To systematically identify regulators associated with fractional resistance, we performed pairwise statistical comparisons of protein expression within the pRB-high population between treated and untreated conditions. Under abemaciclib treatment, many cell cycle regulators (e.g., cyclin D1, p53) showed similar distributions of expression compared to untreated cells (Fig. 3B). However, several regulators including p21, cyclin A2, cyclin E1, CDK2, cyclin B1, and pS6 demonstrated significant shifts in protein expression under 50 nM and/or 250 nM abemaciclib treatment. Notably, p21, an endogenous inhibitor of CDK4 (18), was depleted under treatment, indicating effective pharmacologic inhibition of CDK4 even in proliferating cells.

Cyclin A2 and cyclin E1 were both modestly enriched in fractionally resistant cells. In parallel regulators, phosphorylation of ribosomal protein S6 (pS6), a downstream effector of mTOR signaling associated with active protein synthesis and cell growth, also showed significant shifts within the proliferating cells (19). Total pS6 levels were modestly reduced in proliferating cells at 50 nM relative to untreated proliferating cells (−0.17 vs −0.31; *P* < 0.0001). In contrast, pS6 was significantly elevated in the proliferating cells at 250 nM (−0.17 vs 0.64; *P* < 0.0001).

To visualize these observations in fractionally resistant cells, we plotted protein signal intensity by treatment condition separated by pRB/RB status, where pRB-high cells are shown in pink violin plots and pRB-low cells in blue (Fig 3C). CDK2 expression was significantly increased in cycling cells under 250 nM Abemaciclib treatment relative to the cycling cells off treatment (0.39 vs 1.59; P < 0.0001). Notably, cyclin B1 showed markedly stronger enrichment at 250 nM relative to cycling cells off treatment (−0.20 vs 3.44; P < 0.0001) (Fig. 3, B and C). Importantly, these shifts were specific to the pRB/RB-high cells in comparison to the arrested pRB/RB-low cells which did not show comparable induction of CDK2 or cyclin B1 under treatment (Fig. 3C). Representative immunofluorescence images corroborated these population-level trends by showing pRB-positive cells persisting at 250 nM with elevated CDK2 and cyclin B1 signal (Fig. 3D). Together, these findings indicate that fractionally resistant cells occupying the proliferative region are characterized by enrichment of CDK2, cyclin B1, and pS6.

### Fractional resistance in primary liposarcoma cells

To assess the translatability of cell line findings to primary human liposarcoma, we obtained a fresh primary tumor specimen which was dissociated and plated. We then performed similar 24-hour ex vivo treatment and 4i profiling on primary liposarcoma cells. As with the Lipo246 cell line, an empirical cutoff was determined to identify pRB/RB-high (proliferating) and pRB/RB-low (non-proliferating) cells (Fig. 4A).

**Figure 4.**
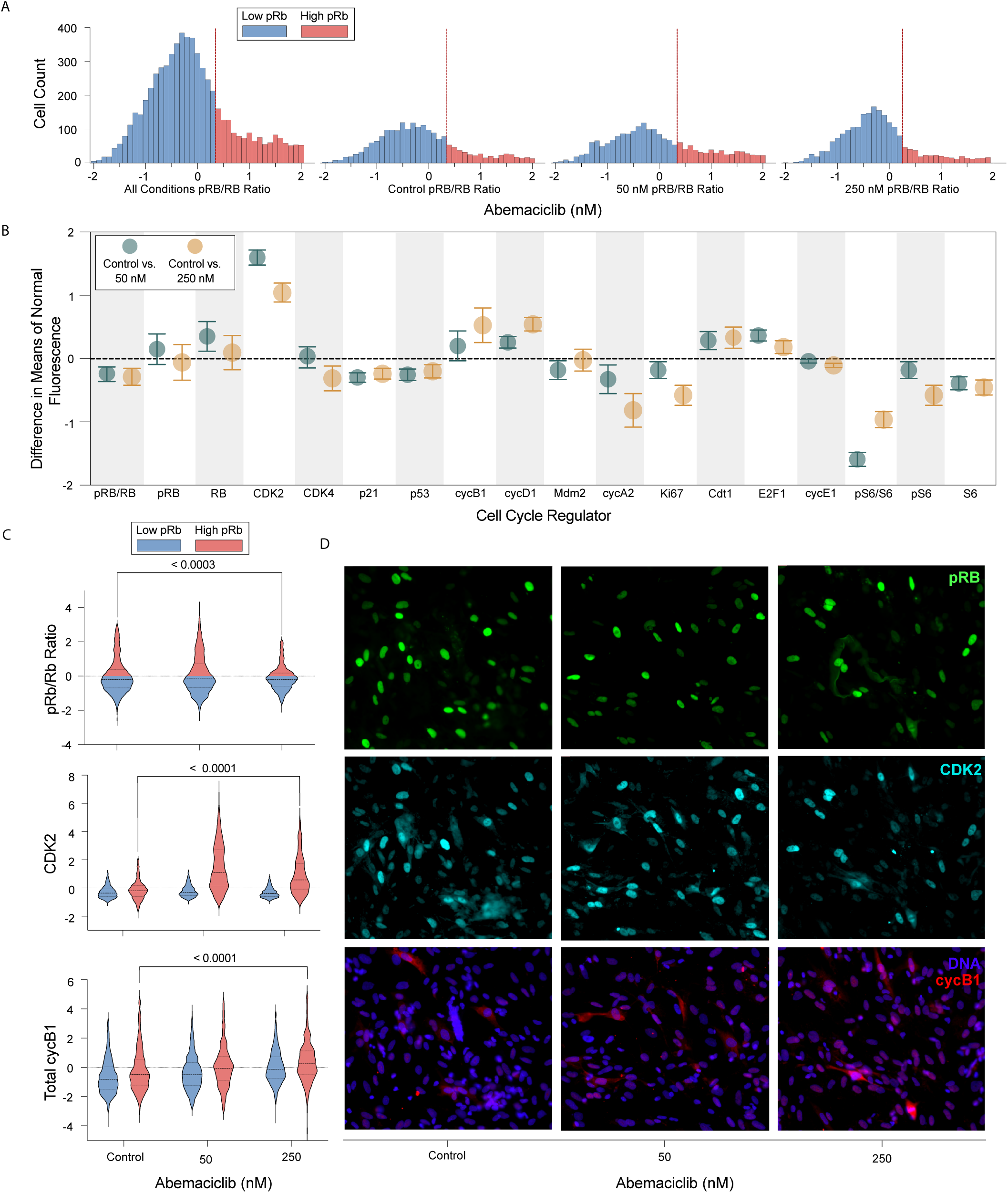
Primary Tumor Cell Cycle Regulator Expression in Fractionally Resistant Cells. (A) Distribution of the ratio of phosphorylated to total RB protein (pRB/RB) across individual primary tumor cells under control (0 nM), 50 nM, and 250 nM abemaciclib treatment. A conservative pRB/RB threshold (vertical red line) was applied to identify cells retaining RB phosphorylation and active proliferation under treatment. (B) Differences in mean expression (normalized z-scores) for cell cycle regulators within the high pRB/RB proliferative fraction, comparing untreated cells with 50 nM (blue) or 250 nM (orange) abemaciclib. Points indicate mean differences and error bars denote 95% confidence intervals; intervals overlapping zero indicate nonsignificant differences. (C) Violin plots showing the distributions of selected cell cycle regulators in low pRB/RB (nonproliferating, blue) and high pRB/RB (proliferating, pink) primary tumor cells across treatment conditions. CDK2 and cyclin B1 (cycB1) exhibit selective enrichment within the proliferative fraction under high dose abemaciclib, consistent with fractional resistance. This enrichment is shown to be statistically significant by ANOVA analysis as seen in the corresponding violin plots. D) Representative images of the effect of dedifferentiated liposarcoma primary tumor cells were treated for 24 hours with 0 (control), 50 nM or 250 nM abemaciclib. Cells were labeled for phosphorylated RB (pRB) which is a marker for actively cycling cells (top panel), cyclin dependent kinase 2 (CDK2)(middle panel) or DAPI for DNA and cycB1 (bottom panel). DAPI staining in the bottom panels shows roughly equal amounts of cells in each drug condition. We can visually observe the colocalization of the upregulation of CDK2 and cycB1 in cells expressing pRB at high abemaciclib dose.

Abemaciclib significantly reduced pRB/RB relative to control at both 50 nM (24.4%) and 250 nM (13.75%) *(P* = 0.0004), demonstrating a functional model with on-target abemaciclib effects. As was seen in the cell line, a proliferating subpopulation of primary tumor cells persisted despite treatment with a high dose of abemaciclib.

We next analyzed features that were enriched in proliferating cells on treatment compared to proliferating cells off treatment to determine features of fractionally resistant cells (Fig. 4B). Generally, the primary cells showed trends similar to the cell line (Fig. 3B). Several regulators exhibited statistically significant but smaller shifts in expression within the proliferating fraction (Fig. 4B). Cyclin D1 increased at both 50 nM and 250 nM (−0.36 vs −0.18 vs −0.055; *P* < 0.0001). Interestingly, for cells cycling under 50 nM treatment, p21 was diminished in comparison to the cycling untreated cells (0.22 to 0.37; Fig. 4B), whereas in the cycling cells under 250 nM, p21 showed a modest increase in comparison to the 50 nM (−0.14 to 0.02; Fig. 4B). The ratio of phosphorylation of ribosomal protein S6 (pS6/S6) showed a similar pattern within the proliferating population. pS6/S6 levels were reduced at 50 nM relative to untreated proliferating cells while pS6/S6 levels in the 250 nM condition had much higher enrichment in comparison to the 50 nM (1.42 vs −0.30 vs 0.41; *P* < 0.0001) (Fig. 4B). By contrast, regulators associated with S-phase entry and replication, including cyclin A2 and cyclin E1, were reduced under treatment. The replication licensing factor Cdt1 showed a slight increase at both 50 nM and 250 nM (−0.23 vs 0.0006 vs 0.10; *P* < 0.0001) (Fig. 4B); however, the magnitude of this effect was modest relative to CDK2 and cyclin B1.

Within the pRB/RB-high proliferative population, CDK2 showed the most pronounced enrichment in fractionally resistant primary tumor cells, under both 50 nM and 250 nM (−0.20 vs 1.10 vs 0.57; *P* < 0.0001) relative to untreated proliferating cells (Fig. 4C). These results indicate persistent accumulation of CDK2 protein in cells that retain RB phosphorylation under CDK4/6 inhibition. In parallel, cyclin B1 was selectively enriched in proliferating cells under a high dose abemaciclib. While cyclin B1 showed only a modest and nonsignificant increase at 50 nM, its expression was significantly elevated at 250 nM (−0.47 vs −0.057 vs −0.26; 250 nM P < 0.0001) (Fig. 4C). Representative immunofluorescence images corroborated these population-level trends by resistant cells persisting at 250 nM with elevated nuclear CDK2 cyclin B1 signal (Fig. 4D). E2F1 was also elevated at 50 nM and 250 nM (−0.20 vs 0.16 vs −0.065; *P* < 0.0001). Importantly, these expression shifts were specific to the pRB/RB-high fraction; pRB/RB-low arrested cells did not show comparable enrichment of CDK2 or cyclin B1 under treatment.

### Co-treatment with a CDK2 inhibitor reveals distinct dose-dependent response behaviors

CDK2 was enriched within the proliferative region under abemaciclib treatment, with the strongest enrichment observed at higher drug doses. Furthermore, canonical CDK2 activating cyclin A2, cyclin E1 and potential noncanonical activating cyclin B1 were significantly enriched in fractionally resistant cells specific to high abemaciclib dose. This dose-specific enrichment suggested fractionally resistant cells may increasingly rely on CDK2-mediated signaling to maintain RB phosphorylation as CDK4/6 activity is suppressed. To test if escaping cells at increasing abemaciclib dose had increased reliance on CDK2 activity, we performed dose response curves to test whether proliferating cells under abemaciclib treatment exhibited altered sensitivity to the CDK2 inhibitor tagtociclib (20). Cells were treated with increasing concentrations of tagtociclib in the presence or absence of abemaciclib, and the proportion of proliferating (high pRB/RB) cells was quantified. Across all conditions, increasing doses of tagtociclib resulted in a dose-dependent reduction in the proliferating population (Fig. 5A).

**Figure 5.**
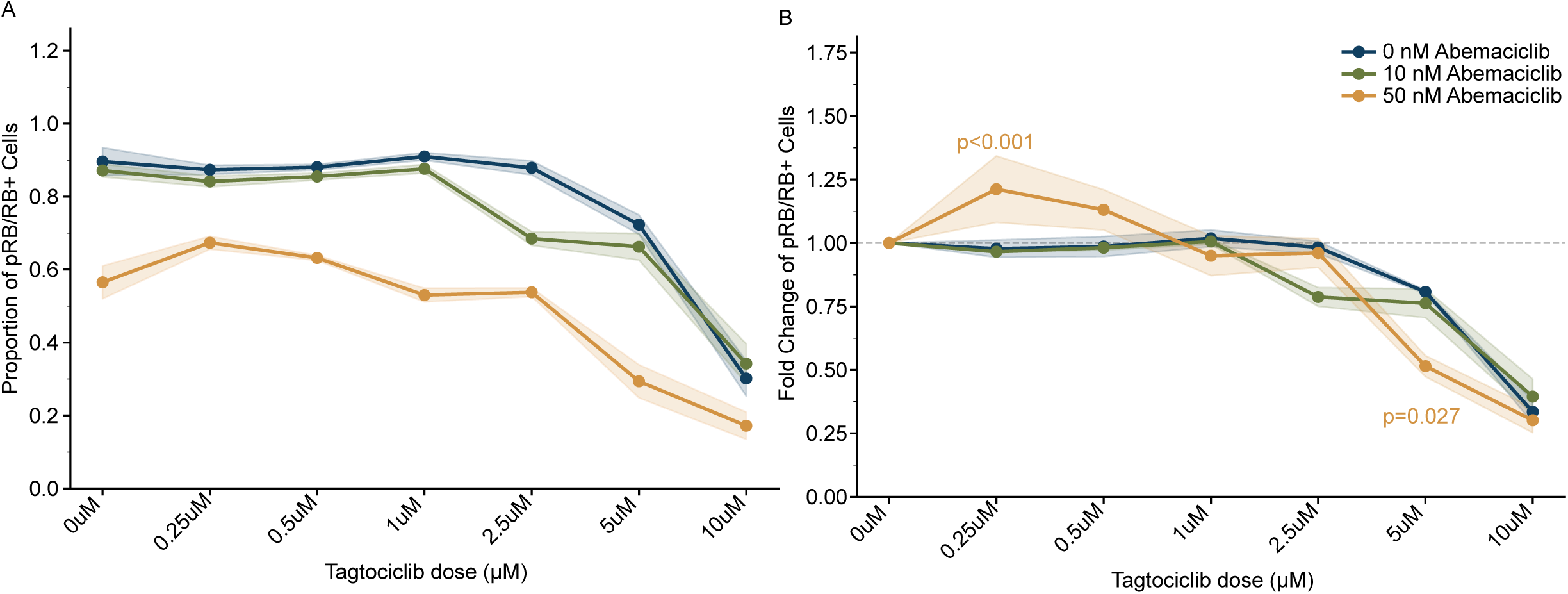
Sensitivity to CDK2 inhibitor tagtociclib at varying doses of abemaciclib. Lipo246 cells were cotreated with the CDK4/6 inhibitor abemaciclib (control, 10 nM, and 50 nM) and a range of CDK2 inhibitor concentrations (0, 0.25, 0.5, 1, 2.5, 5, 10 µM tagtociclib). (A) Proportion of pRB/RB positive cells following treatment with CDK2 inhibitor, tagtociclib. (B) Fold changes in the proportion of pRB/RB positive cells under treatment with tagtociclib. For each abemaciclib condition, fold changes were computed by normalizing the proportion of pRB/RB positive cells to the average proportion of the first three doses of tagtociclib (0, 0.25, and 0.5 µM). Error bars represent the SD across three technical replicates. Dashed lines indicate (A) the proportion or (B) fold change of pRB/RB positive cells without treatment with the CDK2 inhibitor.

To compare dose-dependent interactions between the two treatments, abemaciclib-treated curves were normalized to the corresponding tagtociclib-untreated baseline (Fig. 5B). Across the dose range, two distinct response patterns were observed. At lower concentrations of tagtociclib, increasing abemaciclib dose was associated with maintenance of the high pRB/RB population. In contrast, at higher concentrations of tagtociclib, increasing abemaciclib dose was associated with a greater reduction in the proliferating fraction.

Interestingly, this shift mirrored the dose-dependent enrichment patterns of CDK2 and cyclin B1 observed in both the cell line and primary tumor cells, suggesting a greater reliance on engaging a CDK2 signaling pathway under high abemaciclib dose. Conversely, the apparent antagonism between tagtociclib and abemaciclib at low abemaciclib doses may reflect the partial activity against CDK2 previously observed for abemaciclib (21).

## Discussion

Amplification of CDK4 in dedifferentiated liposarcoma presents a potential therapeutic target, with promising early reports of the CDK4/6 inhibitor abemaciclib (8–10, 22). Herein we characterize liposarcoma cells treated with abemaciclib and identify cell cycle regulator proteins that are enriched in cells that are able to overcome abemaciclib mediated arrest. We show conserved features of resistant cells in the Lipo246 cell line and in primary human dedifferentiated liposarcoma cells. These findings demonstrate reproducible features of plasticity in the cell cycle that allow cells to overcome treatment induced arrest.

Interestingly, the mechanisms of cellular escape to overcome CDK4/6 mediated arrest in liposarcoma share similarities to our prior ER+/HER2− breast cancer study, which may indicate conserved biology of resistance in two very different cancers (13). In that study, we identified cyclinE1 and CDK2 as enriched in fractionally resistant cells to palbociclib treatment, and both of these features are associated with worse response in patients with ER+/HER2− breast cancer receiving palbociclib (13, 23). In this study we identified CDK2 as enriched in fractionally resistant cells, but this is associated with cyclin B1 to a greater degree than cyclin E1 which could represent a non-canonical binding partner activating CDK2 and demonstrate how liposarcoma cells adapt the interaction of cell cycle regulators to resist treatment. Non-canonical Cyclin-CDK binding and activation may be a mechanism by which liposarcoma, and potentially other cancer types, overcome cell cycle-inhibiting drugs. Understanding how CDK2 may be activated by cyclin B1 in dedifferentiated liposarcoma cells could provide insight into how tumor cells evade normal cell cycle regulatory circuits as a hallmark of cancer progression.

All patients with dedifferentiated liposarcoma are not uniformly responsive to abemaciclib (10), and intratumoral heterogeneity at the single-cell level generates a fractionally resistant subpopulation that escapes therapy-induced arrest (13). Characterizing the cell cycle regulator expression patterns underlying this heterogeneity may identify features associated with differential sensitivity to abemaciclib. We characterized cells that continue to divide in the presence of abemaciclib to assess the early biologic response, which may identify the biology that underlies the emergence of resistance. We would expect that tumors that express higher levels of genes enriched in resistant cells would be less sensitive to abemaciclib. Currently no dataset exists to test enriched proteins, including cyclin B1 and CDK2, in fractionally resistant cells as biomarkers of response to abemaciclib. The SARC041 phase 3 clinical trial is currently evaluating abemaciclib as monotherapy in unresectable and metastatic dedifferentiated liposarcoma, and future studies should test if cyclin B1 and CDK2 expression are associated with reduced sensitivity to abemaciclib (22).

Understanding the pathways that permit cells to continue to divide despite abemaciclib treatment may identify cotreatment strategies that enhance efficacy. First, as we showed with ER+/HER2− breast cancer cells, CDK2 is enriched in fractionally resistant dedifferentiated liposarcoma cells. However, unlike ER+/HER2−breast cancer cells, a clear pattern of synergy of abemaciclib with CDK2 inhibition was not seen. While CDK2 inhibition reduced the proliferating population across conditions, two distinct patterns of behavior were revealed. At lower levels of CDK2 inhibition, proliferating cells persisted despite abemaciclib treatment, whereas stronger CDK2 inhibition was associated with a more pronounced reduction in the proliferating population. It has been reported that abemaciclib exhibits partial inhibitory activity against CDK2 in addition to CDK4/6 (10, 24). This broader kinase activity may contribute to its increased clinical efficacy relative to more selective CDK4/6 inhibitors but may also result in enrichment of CDK2 levels that are not associated with increased CDK2 activity and therefore do not confer increased sensitivity to CDK2 inhibitors. Supporting this concept, we saw enrichment of the G2 cyclins (cyclin A2, cyclin B1), rather than the predominantly G1 cyclin E1, suggesting that some of the pRB/RB high cells are arrested in G2 rather than actively cycling. As abemaciclib has previously been shown to promote G2-arrest (21), validating methods to identify cells arrested in G2 may be important to precisely identify the fraction of cells that resist abemaciclib mediated arrest. Alternatively, rather than indicating a single dominant dependency, these results are consistent with a model in which tumor cells retain the capacity to adaptively engage multiple regulatory pathways to preserve proliferation.

While we observed dose dependent synergy of abemaciclib and a CDK2 inhibitor *in vitro*, achieving this window of synergistic concentrations at the tumor in patients may be challenging. We did identify an additional promising target. Enrichment of pS6 in fractionally resistant cells indicates increased activity of the PI3K-mTOR pathway. Importantly, co-treatment with a PI3K inhibitor in addition to the CDK4/6 inhibitor palbociclib has been shown to improve outcomes in PIK3CA mutant ER+/HER2− breast cancer (25, 26), and the VIKTORIA-1 trial showed similar findings in PIK3CA wild-type ER+ tumors (27). Dedifferentiated liposarcoma rarely harbor PIK3CA mutations (28) but may activate this pathway in response to CDK4/6 inhibition. Further work is needed to determine how PI3K pathway activity may facilitate escape from abemaciclib mediated arrest, and the efficacy of targeting this pathway to enhance response rates.

By leveraging single-cell, high-dimensional profiling of both established cell lines and primary human tumor samples, we resolve intratumoral heterogeneity at the level of individual cells and apply computational tools to characterize abemaciclib resistant dedifferentiated liposarcoma cells. The ability to identify and characterize these fractionally resistant populations provides a framework for understanding early adaptive responses to therapy and for developing biomarkers that more accurately predict patient responses. Extending these approaches across tumor types may facilitate the development of more effective, personalized therapeutic regimens aimed at eliminating proliferative escape routes and improving clinical outcomes in patients with aggressive solid tumors.

## Methods

### Sex as a biological variable

Sex was not considered as a biological variable in this study. The Lipo246 cell line and the primary tumor specimen were analyzed for cell-intrinsic responses to CDK4/6 and CDK2 inhibition, and the findings are expected to be relevant across sexes.

### Cell Line Lipo246

The cell line Lipo246 (RRID:CVCL_4U73) was used as an in vitro model of dedifferentiated liposarcoma. This human cell line was originally derived from a retroperitoneal dedifferentiated liposarcoma and has been previously described as a model for studying therapeutic resistance in liposarcoma (10). Cells were grown using Dulbecco modified Eagle medium (DMEM, Gibco) with 10% fetal bovine serum (FBS), 1% Penicillin-Streptomycin at 37°C in a humidified incubator with 5% CO₂. For CDK4/6 inhibition experiments, cells were allowed to adhere on Poly-D-lysine (Gibco) and Fibronectin (Sigma) coated plates at approximately 12,000 cells per well for 24h before treatment with a CDK4/6 inhibitor. Abemaciclib (Selleckchem, catalog no. S5716) was reconstituted in dimethyl sulfoxide (DMSO) and diluted in culture medium to a final DMSO concentration of 0.1%; vehicle-treated controls received 0.1% DMSO in culture medium. Cells were treated with increasing concentrations of abemaciclib (10, 25, 50, 100, 250, and 500 nM) with an IC50 calculated to be 74 nM. Comparative analyses focused on untreated control, 50 nM, and 250 nM treatment conditions. After treatment, cells were fixed with paraformaldehyde and processed for iterative indirect immunofluorescence imaging (4i) (Fig. 1) (14).

### CDK4/6 and CDK2 Cotreatment Lipo246

To evaluate sensitivity to combined CDK4/6 and CDK2 inhibition, Lipo246 cells were cotreated with abemaciclib (Selleckchem, catalog no. S5716) and the CDK2 inhibitor tagtociclib (MedChem Express, catalog no. HY-137894A). Both compounds were reconstituted in DMSO and diluted in culture medium to a final DMSO concentration of 0.1%; vehicle-treated controls received 0.1% DMSO in culture medium. Cells were cotreated with abemaciclib (vehicle control, 10 nM, or 50 nM) and a range of tagtociclib concentrations (0, 0.25, 0.5, 1, 2.5, 5, and 10 µM) for 24 hours. Three technical replicates were performed per condition. Following treatment, cells were fixed with paraformaldehyde and processed for iterative indirect immunofluorescence imaging (4i) as described below.

### Primary Tumor Cells

Primary tumor cells were obtained from a patient with dedifferentiated liposarcoma resected from the retroperitoneum under an Institutional Review Board–approved protocol at the University of North Carolina at Chapel Hill. Written informed consent was obtained and deidentified clinical data were accessed through an institutional honest broker system.

The tumor specimen was obtained in the operating room suite immediately following surgical resection and transferred to the laboratory. Fresh tumor tissue was placed in Dulbecco modified Eagle medium (DMEM, Gibco) with 10% fetal bovine serum (FBS), 1% Penicillin-Streptomycin and transported on ice. The specimen was sharply minced into 2–4 mm fragments prior to enzymatic dissociation. Tumor dissociation was performed using Gentle Collagenase/Hyaluronidase (Stemcell Technologies, catalog no. 07919) in DMEM/F12 supplemented with 5% bovine serum albumin, hydrocortisone, HEPES buffer, and GlutaMAX overnight at 37 °C with agitation. Cells were then centrifuged and washed twice with phosphate-buffered saline supplemented with fetal bovine serum and HEPES buffer. Red blood cells were removed by incubation in ammonium chloride solution (Stemcell Technologies, catalog no. 07800). Cells were subsequently centrifuged and briefly treated with warm 0.05% trypsin-EDTA with DNase I to generate a single-cell suspension. Cells were washed and resuspended in DMEM with 10% FBS. The suspension was sequentially filtered through 100 µm and 40 µm cell strainers to remove debris and large aggregates. Viable cells were counted using trypan blue exclusion and a Countess automated cell counter (Life Technologies). Cells were allowed to adhere on Poly-D-lysine (Gibco) and Fibronectin (Sigma) coated plates at approximately 12500 cells per well for 24h before treatment with a CDK4/6 inhibitor. Abemaciclib was reconstituted in DMSO and diluted in culture medium to a final DMSO concentration of 0.1%; vehicle-treated controls received 0.1% DMSO in culture medium. Cells were treated with increasing concentrations of abemaciclib (10, 25, 50, 100, 250, and 500 nM). Comparative analyses focused on untreated control, 50 nM, and 250 nM treatment conditions. After treatment, cells were fixed with paraformaldehyde and processed for iterative indirect immunofluorescence imaging (4i) as described below (14).

### Iterative Indirect Immunofluorescence Imaging (4i)

Single-cell proteomic measurements for samples were obtained and quantified using iterative indirect immunofluorescence imaging (4i) by adapting protocol of Gut et al. (14). Samples were rinsed three times with phosphate-buffered saline (PBS; pH 7.4) and permeabilized with 0.1% Triton X-100 for 15 minutes at room temperature. Nuclear DNA was stained using Hoechst (1:2500 dilution in PBS) for 15 minutes. Following PBS rinse, wells were imaged to confirm cell distribution and suitability for 4i prior to antibody labeling.

Prior to the first staining round, samples were eluted to improve antibody accessibility. Wells were rinsed three times with distilled water and incubated with elution buffer composed of L-glycine (0.5 M), urea (3 M), and guanidinium chloride (3 M) supplemented with TCEP-HCl (70 mM) and HCl to a pH of 2.5. Cells were washed with elution buffer for three rounds of 10 minutes each with gentle agitation, followed by PBS rinse.

Cells were then incubated for 1 hour at room temperature in 4i blocking solution consisting of maleimide (100 mM) and ammonium chloride (100 mM) added to conventional blocking solution (1% bovine serum albumin in PBS). Primary antibodies diluted in conventional blocking solution were applied at 50 µL per well and incubated overnight at 4°C with gentle agitation. Primary antibodies were selected from distinct host species to permit multiplex detection. After incubation with primary antibodies, wells were rinsed once with PBS and washed three times for 5 minutes each.

Fluorescent secondary antibodies directed against the host species of the primary antibodies were then applied at a dilution of 1:500 in conventional blocking solution together with Hoechst nuclear stain (1:2500). Secondary antibody solutions were incubated for 1 hour at room temperature under light-protected conditions. The secondary antibodies used were donkey anti-rabbit 488, donkey anti-mouse 568, and donkey anti-goat 647. Cells were washed in PBS as described above prior to imaging.

For imaging, wells were filled with imaging buffer consisting of N-acetylcysteine (700 mM, pH 7.4). Fluorescence images were acquired using a Nikon Ti inverted microscope equipped with a Plan Apo Lambda 20× objective (NA = 0.75) and an Andor Zyla 4.2P sCMOS camera. 6×6 images were acquired and stitched together to generate complete images for each well and each staining round. After imaging, antibodies were removed by repeating the elution procedure described above. Elution control images were acquired periodically to confirm successful removal of antibody signal before subsequent staining cycles. This process of antibody labeling, imaging, and elution was repeated iteratively to generate multiplex single-cell protein profiles for each experimental condition.

### Antibody Panel

**Table 1.** List of primary antibodies utilized in the 4i rounds of imaging along with host species, vendor information, and dilution utilized in the experiment.

| Primary Antibody | Species | Vendor/Cat# | Dilution |
| --- | --- | --- | --- |
| CDK2 | Goat | R&D Systems/AF4654 | 1:100 |
| CDK4 | Rabbit | Abcam/ ab108355 | 1:400 |
| p53 | Mouse | CST/48818 | 1:400 |
| Mdm2 | Mouse | Abcam/ab16895 | 1:200 |
| Cyclin A2 | Goat | R&D Systems/AF5999 | 1:400 |
| Cyclin B1 | Goat | R&D Systems/AF6000 | 1:400 |
| Cyclin D1 | Rabbit | CST/55506 | 1:200 |
| Cyclin E1 | Mouse | CST/4129 | 1:200 |
| E2F1 | Mouse | Santa Cruz/sc-251 | 1:100 |
| Cdt1 | Rabbit | CST/8064 | 1:200 |
| Ki67 | Rabbit | Abcam/ab15580 | 1:800 |
| p16 | Mouse | Roche/705-4793 | No Dil. |
| p21 | Goat | R&D Systems/AF1047 | 1:200 |
| pS6 | Rabbit | CST/4858 | 1:200 |
| S6 | Mouse | CST/2317 | 1:100 |
| Phospho-Retinoblastoma (pRB) | Rabbit | CST/8516 | 1:1000 |
| Retinoblastoma (RB) | Mouse | CST/9309 | 1:400 |

### Image Processing and Single-Cell Quantification

Raw images were processed using Python. Images from all rounds were aligned so that the same cell could be analyzed across the entire 4i panel. Cell segmentation was performed using Cellpose (29). After segmentation, images were manually reviewed in Napari and masking was performed to remove debris, smears, and noncellular artifacts that could confound signal quantification. The resulting processed data were used to generate single-cell data frames containing protein expression and cell feature measurements for each cell. Raw image processing and downstream single-cell analysis workflows were implemented in Python and adapted from previously published pipelines for multiplexed imaging and fractional resistance analysis, including publicly available code repositories (13, 14, 29, 30, 31).

### Downsampling and Computational Analysis

To enable balanced comparison across treatment conditions, representative downsampling was performed using kernel herding sketching (30). Equal numbers of cells were selected from each treatment condition, with 2,000 representative cells per condition used for downstream analysis. This yielded a final dataset of 12,000 Lipo246 cells across untreated, 10, 25, 50, 100, 250, and 500 nM abemaciclib conditions. Primary tumor datasets were similarly downsampled to 2,000 cells per condition for direct comparison.

### Identification of Proliferating Cells

The ratio of phosphorylated to total RB protein (pRB/RB) was used to quantify proliferative state at the single-cell level. pRB/RB showed a bimodal distribution, with low values corresponding to hypophosphorylated RB in arrested cells and high values corresponding to hyperphosphorylated RB in actively cycling cells (13). A conservative cutoff was applied to isolate the high pRB/RB subpopulation for analysis of proliferating and fractionally resistant cells (Fig. 1C).

### PHATE Dimensionality Reduction

To visualize high-dimensional single-cell proteomic states, nonlinear dimensionality reduction was performed using Potential of Heat-diffusion for Affinity-based Transition Embedding (PHATE) (15). PHATE was applied to the normalized 4i dataset with parameters knn = 100 and t = 7 to generate a low-dimensional representation in which cells with similar protein-expression profiles were positioned proximally. Protein expression and cellular features from 4i, including pRB, Ki-67, CDK2, cyclin B1, and nuclear area, were overlaid onto PHATE embeddings to identify and interpret proliferative cell cycle regions and visualize the distribution of fractionally resistant cells.

### Statistical Analysis

Comparisons of single-cell protein expression between untreated and abemaciclib-treated proliferating cells were performed using two-sample *t-*tests. Mean differences in normalized fluorescence were calculated for each marker and displayed with 95% confidence intervals for untreated versus 50 nM and untreated versus 250 nM conditions. Violin plots were generated to visualize the distribution of normalized single-cell fluorescence values across treatment groups. Statistical significance was defined as *P* < 0.05.

## Supporting information

Supporting Data Values for Figures

## Data Availability

Values underlying graphed data and reported means presented in the main text are provided in the Supporting Data Values file. Tabular data frames of preprocessed single-cell proteomic data for the cell lines and primary tumor samples across doses have been deposited in Zenodo in AnnData format (https://doi.org/10.5281/zenodo.20187010). Additionally, an interactive 3D PHATE viewer is hosted at https://laelise.github.io/ddlps-fractional-resistance/.

## Author Contributions

L.E.B., S.C.W., P.M.S., and J.E.P. conceptualized the project and designed the research; S.C.W. established experimental workflow, performed experiments and preliminary computational analysis. L.E.B. performed experiments, conducted computational analyses, and wrote the manuscript; L.E.B., T.M.Z., and G.A.S. developed analytic tools; L.E.B., S.C.W., T.M.Z., P.M.S., and J.E.P. interpreted data; J.D.B. and E.K.T. contributed to early project development; A.A.W., E.K.T., and H.R. provided technical and material support; L.E.B., P.M.S., J.E.P., T.M.Z., G.A.S., E.K.T., and S.C.W. edited and revised the manuscript; and all authors reviewed and approved the final manuscript. L.E.B. and S.C.W. contributed equally to this work.

## Acknowledgements

The authors sincerely thank the patients who donated samples for this research. The authors also acknowledge support from the University of North Carolina Postbaccalaureate Research Initiative in Science and Medicine (PRISM) program and thank Ander Naugle for experimental assistance.

